# Which results of the standard test in community weighted mean approach are too optimistic?

**DOI:** 10.1101/349589

**Authors:** David Zelený

## Abstract

**Questions:** Community weighted mean (CWM) approach analyses the relationship species attributes (like traits or Ellenberg-type indicator values) to sample attributes (environmental variables). Recently it has been shown to suffer from inflated Type I error rate if tested by standard parametric or (row-based) permutation test. Results of many published studies are likely influenced, reporting overly optimistic relationships that are in fact merely a numerical artefact. Can we evaluate results of which studies are likely to be influenced and how much?

**Methods:** I suggest that hypotheses commonly tested by CWM approach are classified into three categories, which differ by assumption they make about the link of species composition to either species or sample attributes. I used a set of simulated and one simple real dataset to show how is the inflated Type I error rate influenced by data characteristics.

**Results:** For hypotheses assuming the link of species composition to species attributes, CWM approach with standard test returns correct Type I error rate. However, for the other two categories (assuming link of species composition to sample attributes or not assuming any link) it returns inflated Type I error rate and requires alternative tests to control for it (column-based and max test, respectively). Inflation index is negatively related to the beta diversity of species composition and positively to the strength of species composition-sample attributes relationship and the number of samples in the dataset. Inflation index is also influenced by modifying species composition matrix (by transformation or removal of species). The relationship of CWM with intrinsic species attributes is a case of spurious correlation and can be tested by column-based (modified) permutation test.

**Conclusions:** The concept of three hypothesis categories offers a simple tool to evaluate whether given study reports correct or inflated Type I error rate, and how inflated the rate can be.

## Introduction

A common task of community ecologists is relating species attributes to sample attributes using the matrix of species composition. Species attributes are characteristics of individual species, like species functional traits, ecological optima or phylogenetic age, while sample attributes are characteristics of individual samples, which can be either measured or estimated (environmental variables) or derived from the matrix of species composition itself (species richness, sample ordination scores). The matrix of species composition, which connects species and sample attributes, represents abundances (or presences-absences) of species in community samples. One way to find out whether there is a link between species and sample attributes is to calculate the mean of species attributes for species occurring in each sample weighted by relative species abundances (community-weighted mean, CWM), relate it to sample attributes, e.g. by correlation, and test this relationship by a relevant test. Here I call this method a *CWM approach* and use it as a general term including a wide range of analyses relating CWM of species attributes to sample attributes, where one or several CWMs and one or several sample attributes are involved. The mean of species attributes not weighted by species abundances is also included in CWM approach, since it is identical with CWM calculated on species composition matrix with species presence-absences instead of abundances.

In vegetation ecology, the two commonly used species attributes are plant traits and species indicator values. CWMs of plant traits can be related to environmental variables to demonstrate the effect of environmental filtering on trait-mediated community assembly (Díaz et al. 1998; Shipley 2010), or to predict changes in ecosystem properties (such as biomass production or nutrient cycling; Garnier et al. 2004; Vile et al. 2006), or ecosystem services (like fodder production or maintenance of soil fertility; Díaz et al. 2007). CWMs of species indicator values, like those of Ellenberg et al. (1992) or Landolt (1977) for soil reaction, light, temperature and other factors, are used to estimate habitat conditions from known species composition of vegetation samples. These estimates are often related to soil, light or climatic variables (Ellenberg et al. 1992; Schaffers & Sýkora 2000), used for ecological interpretation of compositional changes in unconstrained ordination (Persson 1981) or ecological differences between groups of samples representing different vegetation types or treatments (Chytrý et al. 2009). Other, more specific examples include relating the community specialization index to environmental variables (Clavero & Brotons 2010; Fajmonová et al. 2013; Carboni et al. 2016), or attempts to verify whether plant biomass can be estimated from tabulated plant heights and species composition as the mean of species heights weighted by their cover in a plot (Axmanová et al. 2012).

The CWM approach is also used in other fields, like biogeography (relating grid-based means of species properties, such as animal body size, to macroclimate or diversity; Hawkins & Diniz-Filho 2006), hydrobiology (relating trophic diatom index based on weighted mean of diatom indicator values to measured water quality parameters to assess its reliability; Kelly & Whitton 1995), or paleoecology (one of the transfer functions used to reconstruct acidification of lakes from fossil diatom assemblages preserved in lake sediments is based on weighted means of diatom optima along the pH gradient; ter Braak & Barendregt 1986; Birks et al. 1990).

The recent paper of Peres-Neto et al. (2017), focused on CWM approach, revealed several surprising facts. First, and perhaps the most important finding, is that standard tests analysing CWM-sample attributes relationship have inflated Type I error rate, returning more optimistic results than is warranted by the data. “Standard tests”, in the meaning used in current paper, include parametric tests like *t*-test for correlation and *F*-test for regression or ANOVA, or permutation tests randomising sample attributes (equivalent to randomising rows in the species composition matrix). Second, CWM correlation without applying weights has rather bad mathematical properties, and dominance of a single species can revert the sign of correlation even if the true trait differences are minimal (see also a dandelion example in Šmilauer & Lepš 2014 and worked example in Appendix S1 of ter Braak et al. 2018). Third, the CWM approach is numerically related to the seemingly different fourth-corner problem (Legendre et al. 1997), which relates species and sample attributes via the species composition matrix without explicitly calculating weighted means of species attributes. Fourth, the ‘max test’ (Cormont et al. 2011), which solves the problem of inflated Type I error rate in the fourth-corner approach (ter Braak et al. 2012), does the same in the CWM approach. The max test undertakes two independent permutation tests, one testing species attributes-species composition link and the other sample attributes-species composition link, and chooses the higher P-value as a result. In conclusion, Peres-Neto et al. (2017) suggested to apply max test in all CWM analyses, and possibly replace the CWM approach with the more efficient fourth-corner approach.

Findings of Peres-Neto et al. (2017) will undoubtedly cause a revolution in the analysis of trait-environment, and generally, species attributes-sample attributes relationship; the max test should be included into the toolbox routinely used methods analysing trait-environment relationship, and fourth-corner approach should be more attention than it had so far. It is also quite relevant to expect that scientific literature using CWM approach with standard tests is flooded by overly optimistic studies reporting significant relationships between various species and sample attributes which in fact are merely an analytical artefact. However, use of CWM approach has a long tradition in ecology, and quite often calculating CWM of species attributes and relating them to sample attributes is practical or required by theory. Many studies defining our current empirical knowledge about the trait-environment relationship or efficiency of Ellenberg-type indicator values have been published, and many studies will use this approach in future. What to do with that? How to recognise whether inflated Type error rate influences the results of certain study or not and if yes, how strong is the influence? Moreover, if CWM approach is used in future studies, is it always necessary to replace the standard tests by the max solution? These are some of the questions I will attempt to answer here.

First, I briefly review the use of CWM approach in vegetation ecology, its conceptual link to other methods analysing the relationship of sample and species attributes via a matrix of species composition, and current knowledge about the problem of inflated Type I error rate. Second, I suggest studies using CWM approach to be classified into one of the three categories, based on underlying assumptions about the link of species or sample attributes to species composition. Two of these categories return inflated Type I error rate in case that CWM approach is tested by the standard test, but only one of these categories requires the use of the max test as the only way to control for correct Type I error rate. Third, I acknowledge that sample attributes are of two types, extrinsic (measured independently of species composition matrix) and intrinsic (derived from species composition matrix), and discuss a special case of CWM correlation with intrinsic species attributes. Finally, I use simulated community data to explore how is the rate of Type I error in standard CWM analysis influenced by data characteristics like beta diversity of species composition matrix, the strength of the link between sample attributes and species composition, and the number of samples in the dataset, and then show the same effect using a real vegetation dataset.

## Theory and Methods

### CWM approach in the context of other methods

Three objects are involved in the calculation of CWM approach: a vector of sample attributes (e, e.g. environmental variables), a matrix of species composition (**L**, abundances or presences-absences of species in samples, with samples as rows and species as columns), and a vector of species attributes (**t**, e.g. species traits); naming convention of variables follows Peres-Neto et al. (2017). CWM of species attributes is calculated as 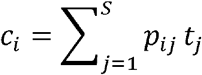, where *S* is the number of species in a community, *p*_ij_ is the relative contribution of species *j* to the total abunance of *i*-th sample, and *t*_*j*_ is the value of species attribute (“trait”) for the species *j*. Relative species proportion *p*_*ij*_ can be calculated as 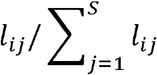, where *l*_*ij*_ is the abundance (or other measure, such as biomass or presence-absence) of species *j* in the *i*-th sample and the denominator is the sum of abundances for all species. The absolute values of *p*_*ij*_ (and consequently also the absolute value of CWM) will be different if the denominator in the formula 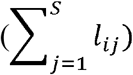 sums species abundances across all species recorded in the data set, or only those for which the values of sample attribute (*t*_*ij*_) is available (and disregarding the others; see more in Discussion). Similar equations (with different notations) are reported in a number of studies, e.g. in Garnier et al. (2004) for CWM of species functional traits or in Diekmann (2003) for CWM of Ellenberg indicator values. CWM is either weighted or unweighted by (absolute or relative) species abundances, which is equivalent to saying that it is calculated on the matrix of species composition using raw (or relative) species abundances (weighted version) or presences-absences (unweighted version). Additionally, CWM can be also weighted by species amplitudes if these are available, where species with narrower amplitudes have a higher weight than species with broader amplitudes. This approach requires extending the CWM formula for amplitude parameter (Zelinka & Marvan 1961), and while commonly used in hydrology (e.g. diatom or saprophytic index, Kelly & Whitton 1995), it is rarely applied in vegetation ecology (but see Peppler-Lisbach 2008 using it for Ellenberg indicator values) and will not be further discussed here.

CWM is related to environmental variables (or other sample attributes) by a wide range of methods like correlation (called CWM correlation in the further text), regression or ANOVA, or more complex methods like mixed effect models, ordination (CWM-RDA and RLQ method, Kleyer et al. 2012; Dolédec et al. 1996) or correlation on distance matrices (Pillar et al. 2009). The strength and the direction of the relationship between CWM and sample attributes are quantified by relevant statistic (model parameters or effect size), and the significance of this statistic is often tested. The test is either parametric (e.g. *t*-test for correlation), or permutation with the test statistic generated by reshuffling sample attributes (equivalent to permuting rows in species composition matrix, hence row-based permutation test, Fig. 1a).

**Fig. 1.**
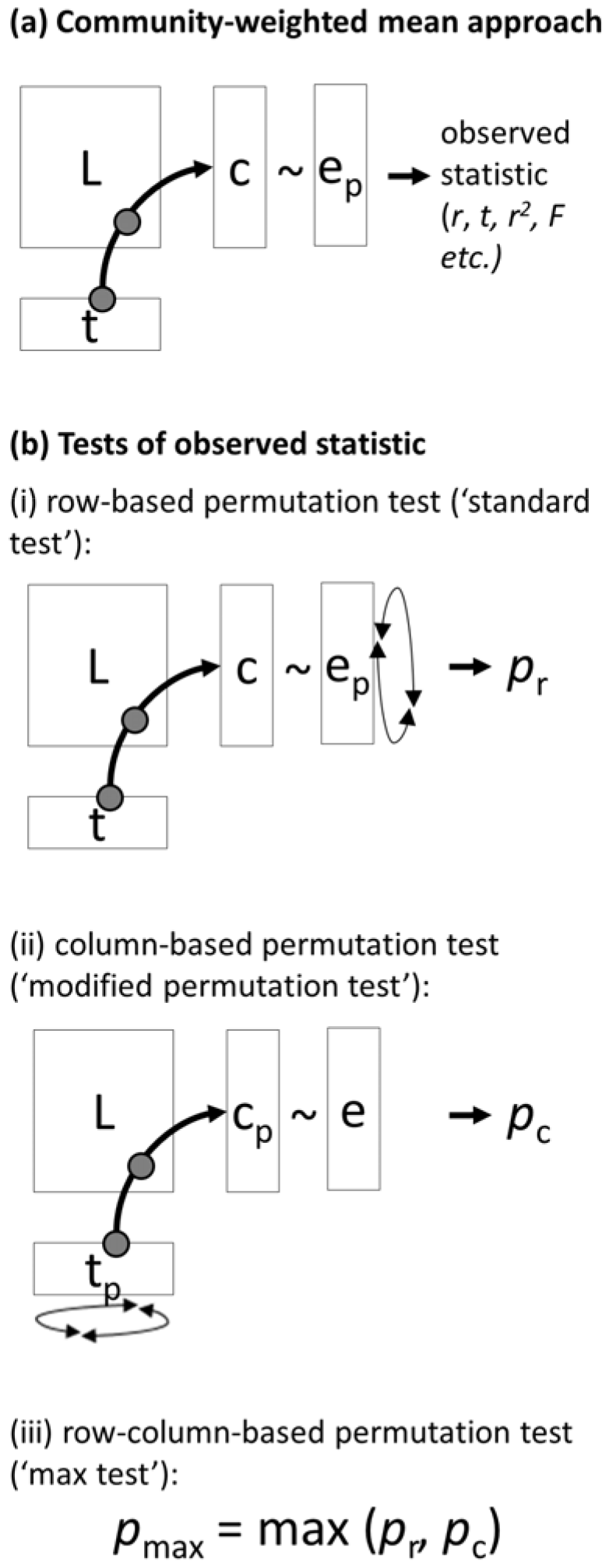
Schema of (a) community-weighted mean approach, which generates the observed value of the test statistic (depending on the method used), and (b) available tests of this statistic. Three tests are available: (i) row-based permutation test (analogy to standard parametric test of c-e relationship), (ii) column-based permutation test (called ‘modified permutation test’ in Zelený & Schaffers 2012), and (iii) max test (called also ‘row-column based permutation test’ in Peres-Neto et al. 2017, or ‘sequential test’ in ter Braak et al. 2012). Notation: **e** = sample attribute (e.g. environmental variable), **t** = species attribute (e.g. trait), **L** = matrix of species composition, **c** = CWM calculated from **t** and **L**, **t**_**p**_ = species attributes permuted among species, **e**_**p**_ = sample attributes permuted among samples, **c**_**p**_ = CWM calculated from **t**_**p**_ and **L**.

Alternative methods analysing pairwise relationships of individual species attributes to individual sample attributes via the matrix of species composition include species niche centroids approach (SNC; ter Braak & Looman 1986) and the fourth-corner approach (Legendre et al. 1997). While in the CWM approach sample attributes are related to the (weighted) mean of species attributes, in SNC approach the species attributes are related to the (weighted) mean of sample attributes (“niche centroids”). The fourth-corner approach (or the “fourth-corner problem”), in contrast, is not explicitly calculating weighted means of species or sample attributes, but combines all three objects (**e**, **t**, and **L**) by inflating the original data tables (Dray & Legendre 2008). The original algorithm by Legendre et al. (1997) considered only presence-absence data in species composition matrix and introduced four different permutation tests, each aiming to test the different ecological hypothesis. Dray & Legendre (2008) extended the method also to quantitative species composition data and introduced universal two-step permutation test, later replaced by max test (ter Braak et al. 2012).

The fourth-corner problem, CWM and SNC approaches are in fact mathematically closely related (Peres-Neto et al. 2017). The fourth-corner statistic *r* is equal to the slope of the weighted linear regression between SNC of environmental variable and trait (Dray & Legendre 2008) or CWM of trait and environmental variable (ter Braak et al. 2018), in the case that the regression is weighted, and traits with environmental variables are weighted standardized prior to calculation. Also, the weighted correlation of CWM of traits and environmental variable or SNC of environmental variable and trait is related to the fourth-corner’s *r*, and can be recalculated to each other using ratios between weighted standard deviations of CWM and traits, or SNC and environmental variable, respectively (equation 15 in Peres-Neto et al. 2017). Weights mentioned above are derived from the species composition matrix **L**, as either total species abundances in samples (row sums in **L**) or sums of individual species abundances across all samples (column sums of **L**). Row sums of **L** are used as weights in CWM regression and correlation and to weighted-standardise environmental variable in the regression, and the column sums of **L** are used as weights in SNC regression and correlation and to weighted-standardise traits in regression. Note that two conceptually different types of weights are mentioned in the context of CWM (and also SNC) method. Abundances of individual species in individual samples (both CWM and SNC can also be calculated unweighted, equivalent to calculating them on matrix of species composition **L** with species presences-absences instead of abundances), and plot or sample weights, calculated as row sums or column sums of species abundances in matrix **L**, respectively (and used as weights in weighted regression and correlation and to weighted standardize environmental variables and traits if necessary). Thus, if using the term “weighted” in case of CWM (or SNC) approach, it is advisable to specify which of the weights are meant.

### Inflated Type I error rate of standard test in CWM approach

As mentioned above, Peres-Neto et al. (2017) showed that CWM approach might return overly optimistic results with inflated Type I error rate, falsely indicating the link between species and sample attributes where there is none. In fact, several previous studies indicated that testing CWM-environment relationship is possibly problematic. Pillar et al. (2009) used column-based permutation test to asses link between CWM and environment using correlation between distance matrices in the study discriminating trait-convergence and divergence patterns in community assembly; they argue that “[t]he null model should retain most of the real data structures except for the one that is to be tested”. Jansen et al. (2011) calculated the relationship between mean Ellenberg indicator values or mean trait values and environmental variables and tested it by randomization test with permutation of species attributes, arguing that “[d]ue to the non-random co-occurrence of species in relevés … the correlation of mean trait values to measured site conditions can also be achieved by chance”. Zelený & Schaffers (2012) warned against the danger of overly optimistic results in the context of relating mean Ellenberg indicator values to ordination scores, assignment of samples into groups using cluster analysis, and species richness. They argued that these optimistic results are caused by CWM inheriting information about the compositional similarity between community samples and relating CWM to other variables having the same similarity issue causes the problem. They suggested that this relationship should either not be tested, or the “modified permutation test” with randomisation of species attributes should be used. Peres-Neto et al. (2012) discussed similar issue in the context of metacommunity phylogenetics, Šmilauer & Lepš (2014, p. 158) in the context of the CWM-RDA method and Hawkins et al. (2017) in the macroecological context when relating CWM of species traits to species richness.

Parallel to developments related to the CWM approach, Dray & Legendre (2008) identified the problem of inflated Type I error rate in the fourth corner (Legendre et al. 1997) if the fourth-corner statistic is tested by the row-based permutation method. Dray & Legendre (2008) suggested the use of two-step testing procedure combining row- and column-based permutation tests together, a method which ter Braak et al. (2012) improved by introducing the sequential testing approach, called also max approach by later studies (ter Braak et al. 2017). The max test, first used by Cormont et al. (2011), is based on taking the maximum *P*-value from sequentially conducted row- and column-based permutation tests.

Hawkins et al. (2017) pointed out an important theoretical difference between *intrinsic* and *extrinsic* sample attributes, which differ in the relationship to the matrix of species composition. In general, *intrinsic* (sample or species) attributes are mathematically derived from the matrix of species composition, while *extrinsic* (sample or species) attributes have no mathematical relationship to it. Examples of intrinsic *sample* attributes include, e.g. species richness or diversity indices, sample ordination scores, sample assignments into clusters by numerical clustering, and also CWM calculated from species attributes and species composition; extrinsic sample attributes include measured or estimated environmental variables or grouping of samples according to experimental design. Intrinsic *species* attributes are also occasionally used (species scores on ordination axes or species optima calculated by weighted mean from species composition matrix) but are not further discussed here. The max test proposed by Peres-Neto et al. (2017) for CWM correlation applies to test the relationship between CWM and extrinsic sample attributes, and ter Braak et al. (2018) made it clear that there is no theoretical justification to use it for testing the relationship of CWM to intrinsic sample attributes. I suggest (in line with Zelený & Schaffers 2012) that test of CWM with intrinsic sample attributes can be done with column-based permutation test (modified permutation test sensu Zelený & Schaffers 2012), if we consider such relationship as an example of spurious correlation (Brett 2004); more about this below (“*Spurious correlation” of CWM with intrinsic sample attributes).*

### Three categories of hypotheses tested by CWM approach

I suggest that each hypothesis tested by the CWM approach fall into one of the three categories (labelled here as A, B or C, see Table 1 for a summary), depending on assumptions it makes about the link of matrix of species composition to either species attributes or sample attributes, respectively (Fig. 2). One may assume that the link exists if there is sufficient support for it either in the explicit formulation of a tested hypothesis or implicitly from the theoretical context of the study. The hypotheses in *category A* assume the link of species attributes to species composition (**t** <-> **L**), hypotheses in *category B* assume the link of sample attributes to species composition (**e**<->**L**), and in hypotheses in *category C* does not assume any of the two links.

**Table 1.** Overview of the characteristics of the three categories of hypotheses tested by the CWM approach. For each category, the corresponding assumption about a link between sample attributes (**e**) or species attributes (**t**) and species composition (**L**) is provided (**x** <-//-> X: no link, **X** <->X: link), as well as the null vs alternative hypothesis and the recommended test.

| Category of hypotheses | A | B | C |
| --- | --- | --- | --- |
| **The assumption about the link between objects (explicit or implicit)** | **t ↔ L** | **e ↔ L** | no assumption |
| **Null hypothesis** | **e $\nleftrightarrow$ L** | **t $\nleftrightarrow$ L** | **e $\nleftrightarrow$ t,**<br>i.e. **e $\nleftrightarrow$ L**<br>and/or **t $\nleftrightarrow$ L** |
| **Alternative hypothesis** | **e ↔ L** | **t ↔ L** | **e ↔ t,**<br>i.e. **e ↔ L** and **t ↔ L** |
| **Recommended test** | **standard parametric test, row-based permutation test** | **a column-based permutation test (modified permutation test)** | **max test** |

**Fig. 2.**
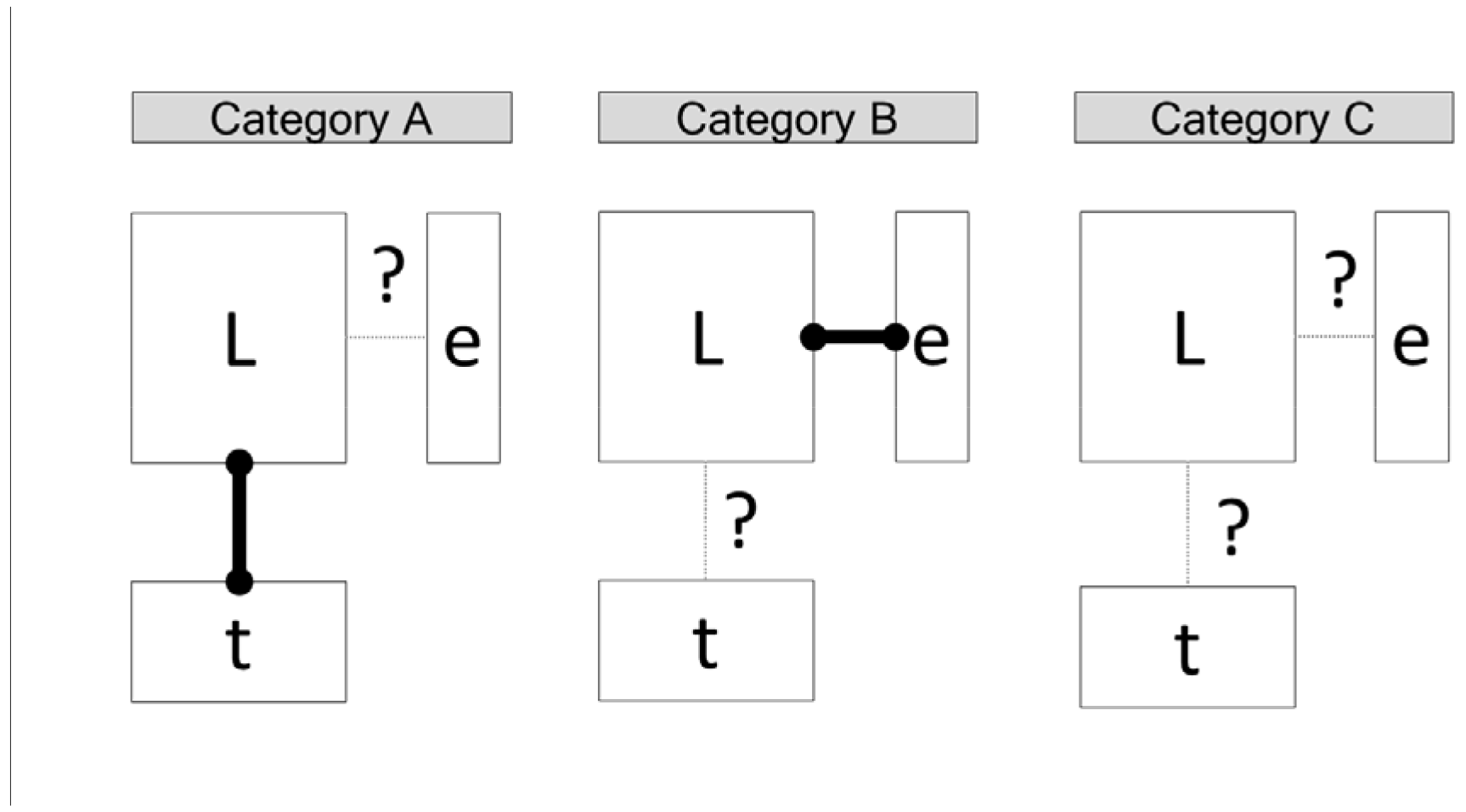
Differences in assumptions behind three categories of hypotheses tested by CWM approach. The bold link indicates that the hypothesis explicitly or implicitly assumes the link between the matrix of species composition (**L**) and either the vector of species attributes (**t**) or vector sample attributes (**e**) and this link is therefore not tested. Question mark, on the other hand, indicates that this link is not explicitly or implicitly acknowledged and can be tested.

Indeed, the choice of the appropriate category may not always be straightforward. For example, trait studies testing whether the environment is filtering the species into a community via their functional traits routinely assume that such traits are functional and as such traits are considered to be linked to species composition (category A). This is reasonable in case that for studied trait there is sufficient evidence from other studies about its functional effect. However, this assumption may not be justified if the analysis is based on traits that are relatively easy to measure and thus readily available in databases, but which may not necessarily be the functional ones. Also, even the trait which is generally considered as functional does not need to be functional in the context of used dataset. Similarly, it may be reasonable to assume that species composition is linked to sample attributes (**L-e**), e.g. if the study is based on experimental treatment which is known to change species composition and the question is focused on how these changes are reflected by sample attributes (e.g. Ellenberg-type indicator values, Chytrý et al. 2009).

As a simple rule to decide whether it is relevant to consider the existence of **L-t** or **L-e** link or not, one may ask whether it is interesting to test the existence of given link, or whether it would make sense to randomise species (**t**) or samples (**e**) attributes, respectively. If the answer is no, it may be safe to assume that given attributes are linked to species composition. In the case of functional traits example above, if there is a sufficient evidence to say that the trait is functional (e.g. experimental study, or previous empirical studies), it may be reasonable to assume that the link exists and does not need to be tested; if any doubt occurs, better to test it. The link between species composition and sample attributes (**L-e**) is tested by row-based test (parametric or permutation), the link between species composition and species attributes (**L-t**) by column-based permutation test, and both links simultaneously by the max test combining both row- and column-based tests together by selecting the larger *P*-value (Peres-Neto et al. 2017 and Fig 1b here). Even if max test seems to represent universal testing solution, in fact in categories A and B, the link which is assumed to exist does not need to be tested. This simplifies the test to either row-based (i.e. standard test), testing the link between sample attributes and species composition in category A, or column-based, testing the link between species attributes and species composition in category B. Only hypotheses in category C require both row- and column-based tests to be done, and max test was proved to control Type I error rate (Peres-Neto et al. 2017).

***Category A.*** Studies in this category assume that species attributes are linked to species composition. For example, trait-based studies asking whether species traits can explain the effect of environmental filtering on species abundance in a community fall into this category. The null hypothesis, which states that sample attributes are not linked to species composition, can be tested by row-based (standard) parametric or permutation test.

***Category B.*** Studies in this category assume that sample attributes are linked to species composition. Includes experimental studies in which the effect of experimental treatment (sample attribute) on species composition is acknowledged, and the question is about the response of species attributes to it. The null hypothesis, which states that species attributes are not linked to species composition, can be tested by column-based permutation test (also called modified permutation test in Zelený & Schaffers 2012).

***Category C.*** Studies in this category assume no link between either species or sample attributes to species composition. Examples include empirical studies describing the general relationship between sample attributes and species attributes, without acknowledging any assumption based on the mechanism of such relationship (e.g. studies relating the CWM of traits to environmental variables without a priori assuming that traits are functional, allowing to question whether particular traits are linked to species composition or not). Studies with species indicator values relating mean indicator values to measured environmental variables also fit this category. To reject the null hypothesis, which states that there is no link between species or sample attributes and the matrix species composition, means to prove that both species and sample attributes are linked to species composition, and this can be done by max test combining both row- and column-based tests.

### “Spurious correlation” of CWM with intrinsic sample attributes

Examples of the relationship of CWM of species attributes with intrinsic sample attributes include analyses testing the relationship between CWM of Ellenberg-like indicator values and ordination scores (Zelený & Schaffers 2012) or CWM of traits with species richness (Hawkins et al. 2017). Wildi (2016) argued that testing such relationship violates the requirement on the independence of tested variables and should not be used, and ter Braak et al. (2018) warns against the use of the max test because it is not justified by the theory behind in this context. I suggest that this relationship can be considered as a case of “spurious correlations”, i.e. a relationship between compound variables calculated from the same parent variables (Pearson 1897). Spurious correlations like X/Z ~ Y/Z, X ~ Y/X or X+Y ~ Y (where X, Y and Z are variables related together) are ubiquitous in ecology, often encountered also in plant trait studies when one routinely calculates and tests the relationship between calculated traits (e.g. between specific leaf area, SLA, and leaf area, LA, where SLA is calculated as ratio between LA and leaf dry weight, LDW: SLA ~ LA = LA/LDW ~ LA). Although opinions on how to deal with spurious correlations differ among researchers (Jackson & Somers 1991), general suggestion is to either avoid analysing relationship between compound variables, or to acknowledge their non-independence by testing their observed relationship (e.g. correlation) against the null expectation which would exist even if the parent variables are generated in random. For this, Jackson & Somers (1991) and Brett (2004) suggested generating such null expectation by a permutation model, permuting the variable occurring only on one side of the equation.

CWM of species attributes and intrinsic sample attributes (like ordination scores or species richness) are both functions of a species composition matrix **L**. CWM can be rewritten as *f*_*1*_/(**t, L**) and intrinsic samples attributes as *f*_2_(**L**), where **t** is the vector of species attributes (Ellenberg-type indicator values, traits) and **L** is the matrix of species composition. In the relationship *f*_*1*_(**t, L**) ~ *f*_*2*_(**L**), the compositional matrix **L** is a parent variable occurring on both sides of the equation, in the same sense as in the spurious correlation. The null expectation of the test statistic can be calculated by permuting the trait values among species as in the modified permutation test suggested by Zelený & Schaffers (2012). Modified test changes the original null hypothesis of *no relationship between CWM of species attributes and intrinsic sample attributes* (i.e. *f*_*1*_(**t, L**) <-//-> *f*_2_(**L**)) into *no relationship between species attributes and intrinsic sample attributes* (i.e. **t** <-//-> *f*_2_(**L**)). In this way, the modified permutation test remains a valid tool to correct for inflated Type I error rate when relating CWM of species attributes to sample attributes (e.g. relating mean Ellenberg-type indicator values to scores of unconstrained ordination, Zelený & Schaffers 2012).

### Dependence of inflated Type I error rate on data characteristics

In this section, I illustrate how is the inflation of Type I error rate in CWM approach dependent on three dataset characteristics: compositional heterogeneity (beta diversity) of the species composition matrix, the strength of the link between sample attributes and species composition (**L-e** link), and the number of samples in the community matrix. For this, I use CWM correlation with standard parametric test and apply it on a number of simulated community datasets. Then, I use the real vegetation dataset with Ellenberg-type indicator values to show how the inflation depends on the strength of the environment-species composition relationship.

#### Design of the simulation study

The algorithm generating simulated community data is an extension of COMPAS model proposed by Minchin (1987). Here, I used model structured by two virtual ecological gradients, which is an extension of one-gradient implementation by Fridley et al. (2007). Along each gradient, a number of unimodal species response curves was generated, where each response curve quantifies the probability with which an individual found in a given gradient location is assigned to given species. Species composition of individual samples was then generated by randomly selecting locations along both gradients and assigning given number of individuals (100) into species according to species probabilities at given gradient location. The first gradient is used to define sample and species attributes (locations of samples equals to sample ‘environmental variable’, while optima of species response curve equals to species ‘trait’), while the second gradient is used to modify the beta diversity of the whole dataset (increasing its length together with proportional increase in the number of species results in increased beta diversity of the species composition matrix). Species niche widths are generated as random numbers of uniform distribution between 500 and 1000 units, independently for each gradient. The effective length of the first gradient was arbitrarily set to 500 units (the true length is 1500 units, but only the range between 500 to 1000 units is populated by samples, to avoid gradient edges with a lower density of response curves). The effective length of the second gradient varied between 500 to 4500 units (also with extra 500 units at each side). As a result, each simulated community data set includes a matrix of sample attributes (‘environmental variable’, **e**), species composition (**L**) and species attributes (‘traits’, **t**), where sample attributes and species attributes are linked to species composition. Because the aim is to show what is the probability that CWM correlation will be significant even if the null hypothesis is true (i.e. species attributes are not related to species composition), species attributes were permuted to remove their link to species composition.

Beta diversity of the dataset was modified by increasing the length of the second gradient. I assumed that 500 units of the second gradient represent one community, i.e. enlarging the second gradient from 500 to 5000 units (by steps of 500 units) generated datasets of increasing beta diversity (with 1 to 9 communities). A dataset with a maximum number of communities was also included (max), in which the data are reshaped in the way that no species are shared among any pair of samples. The strength of the relationship between species composition and sample attributes (**L-e** strength) was manipulated by adding random noise to generated values of sample attributes **e**. I also included one intrinsic sample attribute, mathematically derived from the matrix of species composition (**L**) by an unconstrained ordination (sample scores along the first axis of correspondence analysis calculated on log-transformed species composition data, denoted as CA1). The number of samples was manipulated by increasing the density of locations along both gradients where communities were generated, while keeping the length of the gradients constant; this mimics the real situation of sampling increasing number of community samples within the same range of compositional heterogeneity.

I prepared two scenarios, each with one fixed and two variable data characteristics. In *scenario 1,* the number of samples was fixed (100 samples), while beta diversity and the strength of L-e relationship varied; for each combination of beta diversity (1, 3, 5, 7, 9 and max. communities) and **L-e** strength (0.0, 0.2, 0.4, 0.6, 0.8, 1 and CA1) I generated 1000 datasets. In *scenario 2,* the **L-e** strength was kept fixed (0.6), while the number of samples and beta diversity varied; for each combination of sample size (25×2^n^ samples with *n* = {0, 1, 2, 3, 4, 5} and the same levels of beta diversities as in scenario 1) I generated 1000 datasets. For each dataset, I related CWM of t (weighted by species abundances) with e by Pearson’s *r* correlation and tested the significance by standard parametric *t*-test, and then by permutation max test (199 permutations, with absolute *t*-value as a test statistic). I quantified the inflation of Type I error rate in CWM correlation for each combination of data characteristics by inflation index (sensu Lennon 2000) calculated as I(α) = N_o_/N_e_, where α is the nominal significance level, N_o_ is the number of ‘observed’ correlations significant at α level, and N_e_ is the number of ‘expected’ correlation significant at α level (calculated as N_e_ = αN_total_, where N_total_ is the total number of tests). Inflation index quantifies how many times more likely we are to find significant result compared to the test with correct Type I error rate; test with inflation index close to unity has correct Type I error rate. I plotted the inflation index I (α = 0.05) against beta diversity and the strength of **L-e** link (scenario 1) or a number of samples and beta diversity (scenario 2).

#### Design of real data study

Example dataset of real data contain forest vegetation plots sampled by me on the slopes of deep valley of river Vltava, Czech Republic (Zelený & Chytrý 2007). The total of 97 plots of 10×15 m were sampled at even distances along the transect running along the valley slope. All vascular plant species were recorded and their cover estimated using Braun-Blanquet scale (Westhoff & van der Maarel 1978). A subset of 11 environmental variables measured or estimated for each plot was selected (details in Zelený & Chytrý 2007). Species attributes used in this analysis are Czech indicator values for light, temperature, moisture, reaction and nutrients, which are Ellenberg-type species indicator values recalibrated for territory of the Czech Republic (Chytrý et al. 2018). For the analysis presented here, species composition data include only herbs sampled in the forest understory, since indicator values for light are defined only for herbs and juveniles of woody species. Only species that have all five indicator values available were included, others were removed from both species composition matrix and matrix of indicator values; this is to guarantee that all calculated CWM are based on species composition datasets with identical beta diversity. Additionally to 11 extrinsic (measured or estimated) environmental variables, I included also one intrinsic variable, scores of samples along the first axis of correspondence analysis calculated on (log transformed) species composition data. As a result, three matrices were used for CWM correlation: environmental variables (97 samples × 12 variables), species composition (97 samples × 103 species) and Czech indicator values (103 species × 5 indicator values). CWM was calculated as species mean weighted by estimated species abundances transformed into the percentage scale. The strength of **L-e** link for each environmental variable was quantified as variance (R^2^CCA) this variable explains in canonical correspondence analysis (CCA) on log-transformed species composition data, rescaled to maximum variance one explanatory variable could theoretically explain (equal to variance represented by the first axis of correspondence analysis calculated on same species composition data; Šmilauer & Lepš 2014). The use of CCA is inspired by link between its unconstrained version (CA) and fourth-corner approach (Peres-Neto et al. 2017), and although the ordination based on chi-square distances may not be the best method for CWM correlation (which does not apply species and sample weights in calculation), I use it here as a reasonable proxy. Note that the strength of **L-e** in analysis of simulated and real data is quantified by different methods; in simulated data, the strength is manipulated *post hoc* by adding certain level of noise to values of sample attributes (which by construction has strong **L-e** link), while in real data, the strength is calculated as variance in **L** explained by **e**. Since in real study the species composition data are the same for each combination of **t** and **e**, beta diversity is fixed and R^2^_CCA_ reflects only the **L-e** strength. If the beta diversity was left to vary (e.g. by calculating CWM for species attributes with missing values of **t**), the R^2^_CCA_ would reflect both **L-e** strength and beta diversity of **L**.

All analyses were done in R-project (version 3.5.0, R Foundation for Statistical Computing, Vienna, Austria, https://www.R-project.org/); complete R-script is available in Appendix S1. Simulated data were generated by package *simcom* (Zelený, unpublished), and CWM correlation was calculated by *weimea* (Zelený, unpublished; source code of v. 0.1.10 in Appendix S2, the latest version at https://github.com/zdealveindy/weimea).

## Results

In an analysis based on simulated data, all three data characteristics (beta diversity, the strength of **L-e** link and sample size) influenced the inflation index of CWM correlation tested by standard parametric test (Fig. 3). The inflation index is negatively related to beta diversity and positively to the strength of **L-e** link (Fig. 3a,b). In the case of maximum beta diversity (samples does not share any species) the inflation index is approaching unity for all strengths of **L-e** link. Inflation index is also positively related to the number of samples (Fig. 3c), with the highest inflation index for the most homogeneous community (number of communities = 1); for the most heterogeneous community (maximum beta diversity) the inflation index oscillates around unity. The max test applied on the same data removes the problem of inflated Type I error rate from all combinations of the three data characteristics (returning inflation index close to 1).

**Fig. 3.**
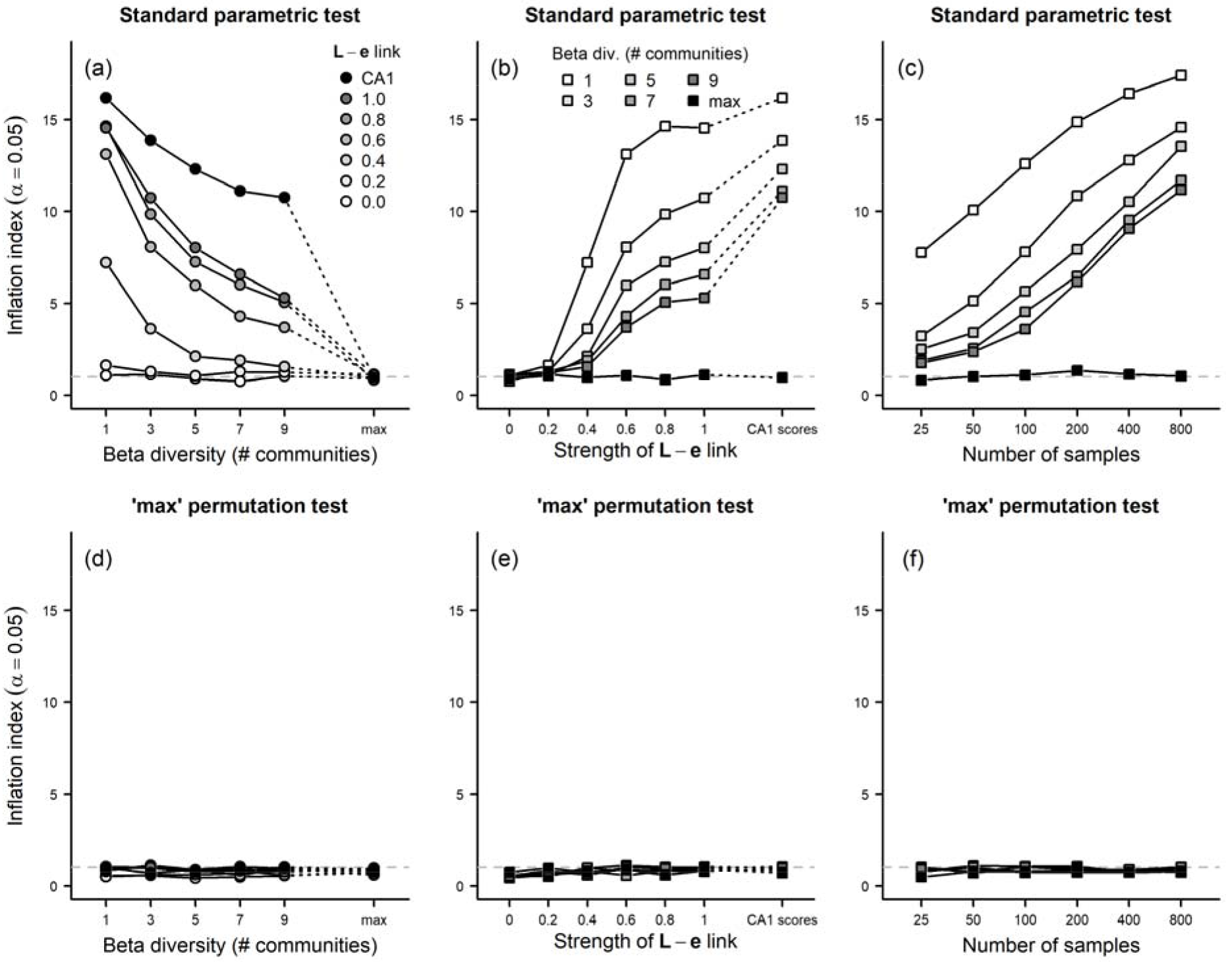
The effect of data characteristics on the inflation index of CWM correlation between CWM of species attributes and sample attributes, tested by parametric t-test of Pearson’s correlation coefficient (panels in top row) and max test (bottom row). Three data characteristics were evaluated: beta diversity of the community dataset (number of communities 1-9 and max. = maximum, when samples in the dataset does not share any species); the strength of the link between sample attributes and species composition (**L-e**; 0 = no link, **e** completely randomized; 1 = full link, generated by the simulation model; CA1 = sample scores on the first CA axis), and the number of samples in the community (25-800). CA1 scores are intrinsic sample attributeswith maximum strength of **L-e** link, since they are derived by correspondence analysis from the community matrix **L**. In each of the panels, one of the characteristics is fixed and the other two are left to vary: in (a, d) and (b, e) beta diversity and L-e link vary, while the number of samples is fixed (n = 100), while in (c, f) the number of samples and beta diversity vary, while the strength of **L-e** is fixed (to value 0.6). The dashed horizontal line is for inflation index equal to one (no inflation).

In the analysis of real data, beta diversity of the species composition data and the number of samples were fixed, and only the strength of **L-e** varied (differ among individual environmental variables). Those environmental variables with a stronger link to species composition were significantly (P < 0.05) related to higher number of CWM of indicator values, with intrinsic variable (CA1) related to all five (Fig. 4a). In an analysis where randomly generated ones replaced real indicator values, the inflation index increased with the strength of L-e relationship, with values over 8 for environmental variables most strongly related to environment and almost 10 for CA1 (Fig. 4b).

**Fig. 4.**
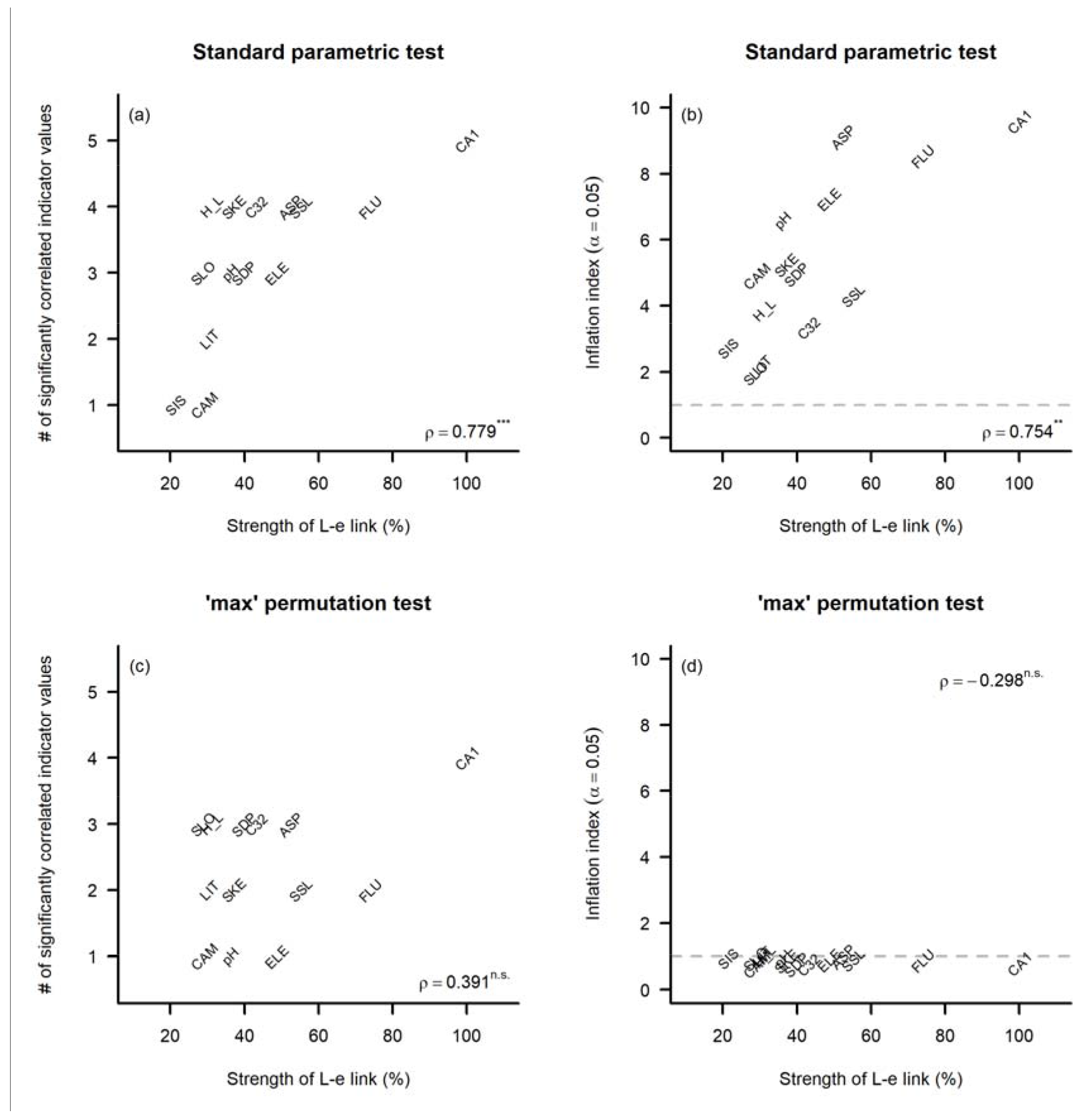
CWM correlations between CWM of Czech indicator values and environmental variables in the real dataset (Vltava), tested by parametric t-test (a, b) and max permutation test (c, d). (a) The number of significant (P < 0.05) correlations between CWM of indicator values and environmental variable (*y*-axis) increases with the strength of **L-e** relationship (*x*-axis, measured as rescaled variance explained by this **e** in CCA). (b) Inflation index of CWM correlation (*y*-axis) also increases with the strength of **L-e** relationship. If the max test replaces standard test, the number of significant indicator values does not relate to the strength of **L-e** relationship (c) and the inflation index is close to unity for all environmental variables (d). ρ = Spearman’s coefficient of correlation between variables on *x-* and *y*-axis (*** = P < 0.001, ** = P < 0.01, n.s. = not significant). Dashed horizontal line (b, d) is for inflation index equal to one (no inflation). Environmental variables: ELE = elevation, SLO = slope, ASP = folded aspect, H_L = heat load, SSL = landform shape in the downslope direction, SIS = landform shape along an isohypse, LIT = presence of lithic leptosols, SKE = presence of skeletic and hyperskeletic leptosols, CAM = presence of cambisol, FLU = presence of fluvisols, SDP = soil depth, pH = soil pH, C32 - cover of tree and shrub canopy.

## Discussion

To avoid inflated Type I error rate in CWM approach, Peres-Neto et al. (2017) suggested using the max test as a universal solution. I suggest that as an alternative to this “one-fits-all” solution, it is useful to fully clarify what are the underlying assumptions the analysed question is putting on the links between members in the game, namely links of species composition to species attributes (**L-t**) or sample attributes (**L-e**), respectively. Standard (row-based parametric or permutation) test controls for Type I error rate for the hypothesis in category A, and results of these studies, therefore, do not need to be considered as overly optimistic. In contrast, hypotheses in categories B and C requires alternative testing approach, namely column-based (modified) permutation test (B) and max test (C), to control for the Type I error rate. This concept can be useful for published studies using CWM approach with standard test, for which one can either clarify whether in the context of given study (with explicitly formulated hypothesis) the standard test returns correct Type I error rate (category A) and if not, whether it is possible to formulate an alternative hypothesis for which the presented results would be valid.

If the study fits the category for which Type I error rate of standard tests is inflated (category B or C), one can evaluate what is the probability that the reported values are overly optimistic. For this, information about beta diversity, the strength of **L-e** relationship and the sample size is needed, or needs to be calculated from the original data (if available). This can also help with conducting a meta-analysis in the future which would evaluate the scale of the problem (how many published studies report overly optimistic results). Indeed, in many published studies the data characteristics are not reported, and original data are not available; then only a rough guess whether the risk is high or low is possible based on available data description. Such guesses are, indeed, only approximate, and re-analysis using the original data is needed to get an exact answer.

Transformation of species abundances (e.g. square-root, log, or presence-absence) will influence the beta diversity of species composition data (and possibly of the strength of **L-e** link) and consequently also inflation index in CWM approach. For example, one may ask whether traits or indicator values are better related to the environment if raw abundances or presences-absences are used for CWM calculation (Hill & Carey 1997; Pakeman et al. 2007). In this sense, the practice differs between the use of traits and indicator values. For traits, the weighting of individual species values by species abundances in the community is justified by Grime’s Mass ratio hypothesis, which states that the functional effect of given species is proportional to its relative contribution to the total biomass of the community (Grime 1998). In contrast, CWM of Ellenberg-type species indicator values more often calculated unweighted by species abundances (i.e. calculated from presence-absence species composition data), because even species with low abundance or biomass can be a good indicator of environmental conditions (Ellenberg et al. 1992). Attempts to answer whether raw abundances or transformed data should be used to calculate CWM are usually done by calculating both CWM of raw and transformed species attributes using the same dataset and relating them to the same sample attributes, including testing the significance by standard tests (Pakeman et al. 2007). This approach, however, does not allow separating the conceptual effect of species data transformation from a mere artefact caused by the fact that data transformation influences inflation index by changes in data characteristics.

Inflation index of standard test in CWM approach is likely to be also influenced by removal of species from species composition matrix, which changes the beta diversity of species composition and strength of L-e relationship. Species are usually removed because they are missing value for given species attribute (e.g. traits measured only for a subset of dominant species, or indicator values without assigning values to generalists) or because of some arbitrary decision (e.g. removing rare species). If more species attributes are related by CWM approach to the same sample attribute using the same species composition matrix, and if these attributes have different proportion and identity of missing species, resulting inflation index can differ among species attributes. This can also bias results of studies that explicitly ask about the sensitivity of CWM approach to missing species values (Ewald 2003; Pakeman & Quested 2007) if these are based on comparing the number of significant relationships of CWM between the same species and sample attributes on datasets with increasing proportion of removed species.

Species with missing values of species attributes values are treated differently in CWM approach applied on traits and Ellenberg-type indicator values. For traits, CWM is often defined as the mean weighted by relative contributions of species into overall biomass, where overall biomass also includes species for which traits were not measured (e.g. Garnier et al. 2004), while for indicator values, species without indicator values (often generalists) are simply disregarded from the calculation (Diekmann 2003). For traits, this is equivalent to calculating species relative contribution *p*_*ij*_ from absolute species abundance divided by abundance sum of all species present in the community (including those with missing trait values). For indicator values, in contrast, species with missing indicator values are first removed from **L** and *p*_*ij*_ is calculated as *l*_*ij*_ divided by sum of *l*_*ij*_ for species left in the matrix. Considering or disregarding the species without species attributes in the CWM calculation changes the absolute value of CWM and in result also the parameters estimated and tested by CWM approach.

In this study, I explicitly ignored intraspecific variation in species attributes, using only dataset-wide mean species attribute values. Indeed, intraspecific variation is important, both in the context of functional traits (Albert et al. 2012) and potentially also Ellenberg-type indicator values (Peppler-Lisbach 2008). In case of traits, intraspecific variability can be considered by calculating CWM values from the site- or treatment-specific species trait values (Lepš et al. 2011). In the case of Ellenberg-type indicator values, species ecological amplitude can be implemented as extra weight in CWM formula (Peppler-Lisbach 2008). Whether and how much is the Type I error rate of such calculations inflated remain to be tested.

Finally, the relevant consideration is whether the CWM approach is the best analytical solution for the question we aim to answer. In cases when the question is explicitly focused on relating community-level values of species attributes, like mean Ellenberg-like species indicator values (serving as an estimate of ecological conditions for individual sites) or the CWM of traits (as one of the functional-diversity metrics and as a community-level trait value) the use of CWM approach is entirely justified. In other cases, when the question is focused on relating individual species-attributes to sample attributes, the fourth corner approach should be considered as it is more powerful in detecting the sample attribute-species attribute relationship (Peres-Neto et al. 2017).

## Conclusions

The CWM approach with standard (row-based) test returns correct Type I error rate only in case the tested hypothesis assume that species composition is linked to species attributes. In other cases, the Type I error rate of the standard test is inflated, and the inflation index depends on the interaction between the beta diversity of species composition matrix, the strength of the relationship between species composition and sample attributes, and the number of samples in the analysis. An alternative to standard test is a column-based or max test, respectively, controlling Type I error rate if species composition is linked to sample attributes (column-based test) or no link is assumed (max test). This concept can be used to evaluate whether results of studies using CWM approach with standard test report correct or inflated Type I error rate, and if inflated, how much.

## Acknowledgements

My thanks go to Cajo ter Braak and two anonymous reviewers for critical comments on previous versions of this manuscript. This study was supported by the Ministry of Science and Technology, Taiwan (106-2621-B-002-003-MY3).

## Supporting information

**Appendix S1.** R-code used to calculate simulated data and real data analysis.

**Appendix S2.** Source code for the R library *weimea,* version 0.1.10 (latest version can be installed from https://github.com/zdealveindy/weimea/).

## Abbreviations

CWM: community weighted mean
SNC: species niche centroid

## References

Albert, C. H., de Bello, F., Lavorel, S. & Thuiller, W. 2012. On the importance of intraspecific variability for the quantification of functional diversity. Oikos 121: 116–126.

Axmanová, I., Tichý, L., Fajmonová, Z., Hájková, P., Hettenbergerová, E., Li, C.-F., Merunková, K., Nejezchlebová, M., Otýpková, Z., Vymazalová, M. & Zelený, D. 2012. Estimation of herbaceous biomass from species composition and cover. Applied Vegetation Science 15: 580–589.

Birks, H. J. B., Line, J.M., Juggins, S., Stevenson, A.C. & ter Braak, C.J.F. 1990. Diatoms and pH reconstruction. Philosophical Transactions of the Royal Society B Biological Sciences 327: 263–278.

Brett M.T. 2004. When is a correlation between non-independent variables “spurious”? Oikos 105:647–656.

Carboni, M., Zelený, D. & Acosta, A. 2016. Measuring ecological specialization along a natural stress gradient using a set of complementary niche breadth indices. Journal of Vegetation Science 27: 892–203.

Chytrý, M., Hejcman, M., Hennekens, S.M. & Schellberg, J. 2009. Changes in vegetation types and Ellenberg indicator values after 65 years of fertilizer application in the Rengen Grassland Experiment, Germany. Applied Vegetation Science 12: 167–176.

Chytrý, M., Tichý, L., Dřevojan, P., Sádlo, J. & Zelený, D. 2018. Ellenberg-type indicator values for the Czech flora. Preslia 90: 83–103.

Clavero, M. & Brotons, L. 2010. Functional homogenization of bird communities along habitat gradients: accounting for niche multidimensionality. Global Ecology and Biogeography 19: 684–696.

Cormont, A., Vos, C.C., van Turnhout, C.A.M., Foppen, R.P.B. & ter Braak, C.J.F. 2011. Using life-history traits to explain bird population responses to changing weather variability. Climate Research 49: 59–71.

Díaz, S., Cabido, M. & Casanoves, F. 1998. Plant functional traits and environmental filters at a regional scale. Journal of Vegetation Science 9: 113–122.

Díaz, S., Lavorel, S., de Bello, F., Quétler, F., Grigulis, K. & Robson, T.M. 2007. Incorporating plant functional diversity effects in ecosystem service assessments. Proceedings of the National Academy of Sciences USA 104: 20684–20689.

Diekmann, M. 2003. Species indicator values as an important tool in applied plant ecology - a review. Basic and Applied Ecology 4: 493–506.

Dolédec, S., Chessel, D., ter Braak, C.J.F. & Champely, S. 1996. Matching species traits to environmental variables: a new three-table ordination method. Environmental and Ecological Statistics 3: 143–166.

Dray, S. & Legendre, P. 2008. Testing the species traits-environment relationships: the fourth-corner problem revisited. Ecology 89: 3400–3412.

Ellenberg, H., Weber, H.E., Düll, R., Wirth, V., Werner, W. & Paulissen, D. 1992. Zeigerwerte von Pflanzen in Mitteleuropa. Second Edition. Scripta Geobotanica 18: 1–248.

Ewald, J. 2003. The sensitivity of Ellenberg indicator values to the completeness of vegetation relevés. Basic and Applied Ecology 4: 507–513.

Fajmonová, Z., Zelený, D., Syrovátka, V., Vončina, G. & Hájek, M. 2013. Distribution of habitat specialists in semi-natural grasslands. Journal of Vegetation Science 24: 616–627.

Fridley, J.D., Vandermast, D.B., Kuppinger, D.M., Manthey, M. & Peet, R.K. 2007. Cooccurrence based assessment of habitat generalists and specialists: a new approach for the measurement of niche width. Journal of Ecology 95: 707–722.

Garnier, E., Cortez, J., Billès, G., Navas, M.L., Roumet, C., Debussche, M., Laurent, G., Blanchard, A., Aubry, D., (…) & Neill, C. 2004. Plant functional markers capture ecosystem properties during secondary succession. Ecology 85: 2630–2637.

Grime, J.P. 1998. Benefits of plant diversity to ecosystems: immediate, filter and founder effects. Journal of Ecology 86: 902–910.

Hawkins, B.A. & Diniz-Filho, J.A.F. 2006. Beyond Rapoport’s rule: evaluating range size patterns of New World birds in a two-dimensional framework. Global Ecology and Biogeography 15: 461–469.

Hawkins, B.A., Leroy, B., Rodríguez, M.A., Singer, A., Vilela, B., Villalobos, F., Wang X. & Zelený D. 2017. Structural bias in aggregated species-level variables driven by repeated species co-occurrences: a pervasive problem in community and assemblage data. Journal of Biogeography 44: 1199–1211.

Hill, M.O. & Carey, P.D. 1997. Prediction of yield in the Rothamsted Park Grass Experiment by Ellenberg indicator values. Journal of Vegetation Science 8: 579–586

Jackson, D.A. & Somers, K.M. 1991. The spectre of ‘spurious’ correlations. Oecologia 86: 147–151.

Jansen, F., Ewald, J. & Zerbe, S. 2011. Ecological preferences of alien plant species in NorthEastern Germany. Biological Invasions 13: 2691–2701.

Kelly, M.G. & Whitton, B.A. 1995. Biological monitoring of eutrophication in rivers. Hydrobiologia 384: 55–67.

Kleyer, M., Dray, S., de Bello, F., Lepš, J., Pakeman, R.J., Strauss, B., Thuiller, W. & Lavorel, S. 2012. Assessing species and community functional responses to environmental gradients: which multivariate methods? Journal of Vegetation Science 23: 805–821.

Landolt, E. 1977. Ökologische Zeigerwerte zur Schweizer Flora. Veröffentlichungen des Geobotanischen Institutes der Eidgenössischen Technischen Hochschule, Stiftung Rübel, Zurich 64:1–208.

Legendre, P., Galzin, R. & Harmelin-Vivien, M.L. 1997. Relating behavior to habitat: solutions to the fourth-corner problem. Ecology 78: 547–562.

Lennon, J.J. 2000. Red-shifts and red herrings in geographical ecology. Ecography 23: 101–113.

Lepš, J., de Bello, F., Šmilauer, P. & Doležal, J. 2011. Community trait response to environment: disentangling species turnover vs intraspecific trait variability effects. Ecography 34: 856–863.

Minchin, P.R. 1987. Simulation of multidimensional community patterns: towards a comprehensive model. Vegetatio 71: 145–156.

Pakeman, J. & Quested, H.M. 2007. Sampling plant functional traits: What proportion of the species need to be measured? Applied Vegetation Science 10: 91–96.

Pakeman, R.J., Garnier, E., Lavorel, S., Ansquer, P., Castro, H., Cruz, P., Doležal, J., Eriksson, O., Freitas, H., …, & Vile, D. 2008. Impact of abundance weighing on the response of seed traits to climate and land use change. Journal of Ecology 96: 355–366.

Pearson, K. 1897. On a form of spurious correlation which may arise when indices are used in the measurement of organs. Proceedings of the Royal Society of London 60: 489–498.

Peppler-Lisbach C. 2008. Using species-environmental amplitudes to predict pH values from vegetation. Journal of Vegetation Science 19: 437–444.

Peres-Neto, P.R., Dray, S. & ter Braak, C.J.F. 2017. Linking trait variation to the environment: critical issues with community-weighted mean correlation resolved by the fourth-corner approach. Ecography 40: 806–816.

Peres-Neto, P.R., Leibold, M.A. & Dray, S. 2012. Assessing the effects of spatial contingency and environmental filtering on metacommunity phylogenetics. Ecology 93: S14–S30.

Persson S. 1981. Ecological indicator values as an aid in interpretation of ordination diagrams. Journal of Ecology 69: 71–84.

Pillar, V.D., Duarte, L.S., Sosinski, E.E. & Joner, F. 2009. Discriminating trait-convergence and trait-divergence assembly patterns in ecological community gradients. Journal of Vegetation Science 20: 334–348.

Schaffers, A.P. & Sýkora, K.V. 2000. Reliability of Ellenberg indicator values for moisture, nitrogen and soil reaction: comparison with field measurements. Journal of Vegetation Science 11: 225–244.

Shipley, B. 2010. From plant traits to vegetation structure. Chance and selection in the assembly of ecological communities. Cambridge University Press, Cambridge, UK.

Šmilauer, P. & Lepš J. 2014. Multivariate analysis of ecological data using CANOCO 5. 2nd ed. Cambridge University Press, Cambridge, UK.

ter Braak, C.J.F, Peres-Neto, P. & Dray, S. 2017. A critical issue in model-based inference for studying trait-based community assembly and a solution. PeerJ 5:e2885.

ter Braak, C.J.F, Peres-Neto, P. & Dray, S. 2018. Simple parametric tests for trait-environment association. Journal of Vegetation Science, accepted.

ter Braak, C.J.F. & Barendregt, L.G. 1986. Weighted averaging of species indicator values: its efficiency in environmental calibration. Mathematical Biosciences 78: 57–72.

ter Braak, C.J.F. & Looman, C.W.N. 1986. Weighted averaging, logistic regression and the Gaussian response model. Vegetatio 65: 3–11.

ter Braak, C.J.F., Cormont, A. & Dray, S. 2012. Improved testing of species traits-environment relationships in the fourth-corner problem. Ecology 93: 1525–1526.

Vile, D., Shipley, B. & Garnier, E. 2006. Ecosystem productivity can be predicted from potential relative growth rate and species abundance. Ecology Letters 9: 1061–1067.

Westhoff, V., & van der Maarel, E. 1978. The Braun-Blanquet approach. In R. H. Whittaker (Ed.), Classification of plant communities (pp. 289–399). W. Junk, The Hague.

Wildi, O. 2016. Why mean indicator values are not biased. Journal of Vegetation Science 27: 40–49.

Zelený, D. & Chytrý, M. 2007. Environmental control of vegetation pattern in deep river valleys of the Bohemian Massif. Preslia, 79: 205–222.

Zelený, D. & Schaffers, A.P. 2012. Too good to be true: pitfalls of using mean Ellenberg indicator values in vegetation analyses. Journal of Vegetation Science 23: 419–431.

Zelinka M. & Marvan P. 1961. Zur Präzisierung der biologischen Klassifikation der Reinheit fliessender Gewässer. Archiv für Hydrobiologie 57: 389–407.

